# Nanoneedles for targeted siRNA silencing of p16 in the Human Corneal Endothelium

**DOI:** 10.1101/2022.05.27.493597

**Authors:** Eleonora Maurizi, Davide Alessandro Martella, Davide Schiroli, Alessia Merra, Salman Ahmad Mustfa, Graziella Pellegrini, Claudio Macaluso, Ciro Chiappini

**Affiliations:** Dentistry Centre Lab, University of Parma, via Gramsci 14, Parma, Italy; Centre for Regenerative Medicine ‘‘S. Ferrari’’, University of Modena and Reggio Emilia, Modena, Italy; Centre for Craniofacial and Regenerative Biology, King’s College London, London, United Kingdom; Holostem Terapie Avanzate S.r.l., Modena, Italy; Transfusion Medicine Unit, Azienda USL-IRCCS, Reggio Emilia, Italy; London Centre for Nanotechnology, King’s College London, London, United Kingdom

## Abstract

Nanoneedles can target nucleic acid transfection to primary cells at tissue interfaces with high efficiency and minimal perturbation. The corneal endothelium is an ideal target for nanoneedle-mediated RNAi aimed at enhancing its proliferative capacity, necessary for tissue regeneration. Here we develop a strategy for siRNA nanoninjection of the human corneal endothelium. We show that nanoneedles can deliver p16-targeting siRNA to primary human corneal endothelial cells *in vitro* without toxicity. The nanoinjection of siRNA induces p16 silencing and increases cell proliferation, as monitored by ki67 expression. Furthermore, siRNA nanoinjection targeting the human corneal endothelium is non-toxic *ex vivo* and silences p16 in transfected cells. These data indicate that nanoinjection can support targeted RNAi therapy for the treatment of endothelial corneal dysfunction.

## Introduction

The cornea is the outermost lens of the eye, our window to overlook the external world by focusing light rays into the eye and allowing vision. Maintaining corneal transparency is essential to guarantee an optimal eyesight, and this is possible only if all the corneal layers are intact and functional. In particular, the inner monolayer, the corneal endothelium (CE), is fundamental for balancing the liquid exchange that guarantees corneal nourishment, clearance and transparency. However, the corneal endothelium has a limited regenerative capacity, as human corneal endothelial cells (HCEnCs) are arrested in the G1 phase of the cell cycle^1^. Therefore, any loss of HCEnCs is permanent, and progressively leads to an impaired liquid exchange across the cornea, which becomes swollen and opaque, causing loss of vision. The only available treatment for diseases affecting corneal endothelial integrity, such as Fuchs dystrophy, ageing or iatrogenic damages, is corneal transplantation, which is the most frequent type of graft performed worldwide^2^. However, corneal transplantation is an invasive procedure, presenting several limitations related to the risk of allogeneic graft rejection and failure, the need for long-term immunosuppressive therapy, and the scarce availability of donor corneas^3^.

Novel approaches for restoring suitable HCEnCs density are key to improving treatment options for corneal endothelial dysfunction. The most appealing and least invasive alternative to corneal transplantation is the regeneration of a patient’s own corneal endothelium through transient induction of HCEnCs proliferation either *in vivo* or *ex vivo*^4^. Moreover, increasing the HCEnCs number through HCEnCs expansion in eye bank corneas would be beneficial to reduce tissue wastage^5^, as the high cell death rate during storage, in particular following apoptosis in the corneal endothelium, leads to a decrease in HCEnCs density^5^ and rejection of more than 35% of stored corneas^2^,^6^.

Gene therapy can increase HCEnCs density in many ways, including by inhibiting apoptosis through overexpression of anti-apoptotic genes (e.g. Bcl-xl)^7^,^8^, by inducing cell proliferation through overexpression of transcription factors (e.g. E2F2)^9^, or by downregulation of cell cycle inhibitors (e.g. p21, p16^10,11^, p27^12^, SNAI1 and CDK2)^13^. Yet, nucleotide delivery to the corneal endothelium is challenging, as the cells are post-mitotic and thus hard to transfect.

Similarly to the retina^14^, the human cornea is an ideal target within the eye for assessing novel gene therapies because of its relative immune privilege and accessibility that implies a minimally invasive surgical manipulation and allows an easy monitoring. Localised delivery is the preferred administration route of gene therapies to the ocular tissues including the corneal endothelium^15^, since the systemic route is not efficient, leading to unfavourable bio-distribution with associated side effects^16^.

Among local gene delivery approaches to the human cornea, viral transduction still raises immunogenicity and safety concerns^17^, steering research towards safer non-viral approaches, using lipid based transfection^*18*^ and electroporation^19,20^. Those non-viral delivery methods can be effective *in vitro* and *ex vivo* but still present cell toxicity, do not provide a localized delivery, and do not efficiently address accessibility challenges *in vivo*.

Nanoneedles are a promising approach for corneal delivery, where conventional topical routes are hampered by a drug bioavailability of around 5% that requires large dosing and frequent administration, with risks of severe side effects^21,22^. Silicon nanoneedles integrated in tear-soluble contact lenses are an efficient and painless solution for long-term delivery of ocular drugs^21^. In particular, the corneal endothelium is an appealing target for nucleic acid nanoinjection^4^. The relative immune privilege of the cornea would reduce the risk of inflammatory response and, since nanoneedles are designed for delivery limited to the superficial layers of a tissue, nanoinjection would selectively reach cells within the corneal endothelial monolayer. Nanoneedle-mediated delivery, known as nanoinjection, efficiently transfects other post-mitotic human cells with high efficiency, without toxicity and with minimal perturbation of cell phenotype^23 24^. In particular, the porous structure of silicon nanoneedles, entirely biodegradable and capable of hosting large payloads, have emerged as a biocompatible platform that efficiently interfaces with living organisms and human tissue for localized gene therapy and molecular diagnostics, with no off target effects^25^.

Here we use porous silicon nanoneedles to develop a nanoinjection approach for RNAi therapy targeting the human corneal endothelium, aimed at restoring HCEnCs proliferative capacity through p16 (CDKN2A) silencing. In this approach, *in vitro* nanoinjection of siRNA targeting p16 into primary human corneal endothelial cells preserves their viability and morphological phenotype, while silencing p16 expression, reducing levels of p16 protein and promoting cell proliferation. Furthermore, nanoinjection targeting the endothelial layer of explanted human corneas preserves cellular structure and does not induce apoptosis while silencing p16 in transfected cells. These results suggest that nanoinjection is a nontoxic method for nucleic acid transfection targeted to the human corneal endothelium.

## Results

### Nanoneedle interfacing with HCEnCs in vitro

We first determined the impact of nanoinjection in primary HCEnCs *in vitro*. Nanoinjection uses a nanoneedle chip loaded with siRNA and placed over the culture, with the nanoneedles facing the cells (Figure 1A). Centrifugation is applied to the system to assist the interfacing. During nanoinjection, confocal microscopy shows multiple nanoneedles co-localising with the cytosol and nucleus of each cell, indicating successful interfacing (Figure 1B). Comparing treated and untreated cells (ctr) on removal of the nanoneedles 30 min following centrifugation, the cells retained their characteristic morphology in culture (Figure1C). The nanoneedle-treated HCEnCs also showed a native ZO-1 pattern, indicating a sealed monolayer that preserves the correct morphology (Figure 1D). The expression of ZO-1 in HCEnCs plasma membranes reveals their characteristic belt of tight junctions, which is strictly connected to their function as a semi-permeable barrier, allowing the regulated diffusion of nutrients from the anterior chamber to the whole cornea^26^. Lack of Caspase3/7 activation revealed that apoptosis in HCEnCs is entirely absent upon nanoinjection, similarly to untreated cells (Figure 1E).

**Figure 1.**
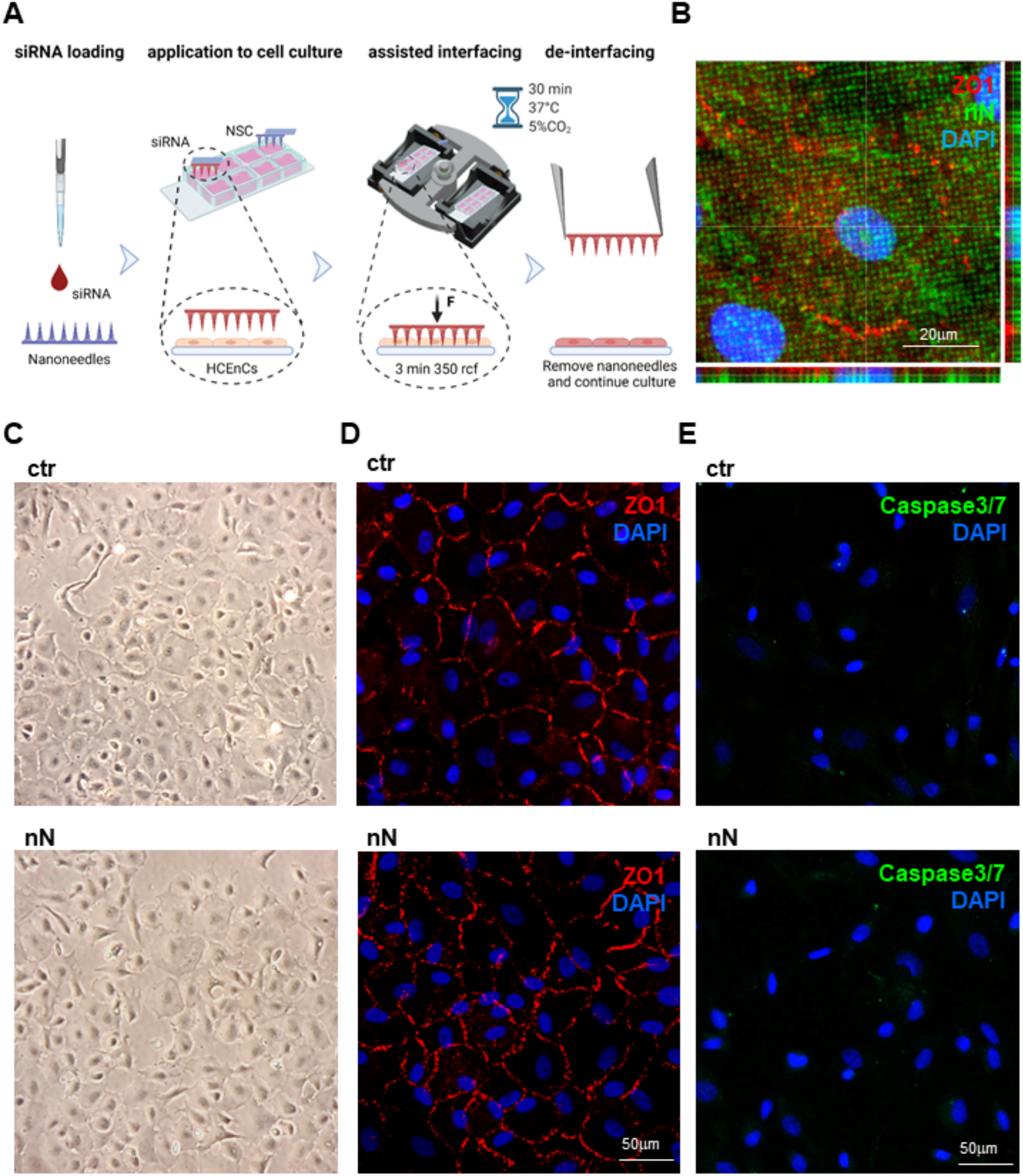
Nanoneedle interfacing with human corneal endothelial cells *in vitro*. (A) Schematic representation of the nanoinjection approach for cultured primary HCEnCs. Image created with Biorender.com (B) Confocal microscopy orthogonal projections of nanoneedles (FITC labelled, green) interfaced with the cytosol, outlined by ZO-1, and the nucleus of HCEnCs. Nanoneedles co-localise with HCEnCs. ZO-1 staining (red) with DAPI (blue) nuclear counterstain. Scale bar 20μm. Images were obtained immediately after nanoneedle assisted interfacing by centrifugation. (C) Phase-contrast image of the primary HCEnCs culture showing retained morphology following nanoneedle interfacing (nN), similarly to the untreated control (ctr). Images were obtained immediately after the de-interfacing. (D) Immunofluorescence microscopy of HCEnCs showing retained hexagonal morphology and ZO-1 marker upon nN interfacing (nN) as well as in untreated HCEnCs (ctr). ZO-1 staining (red) with DAPI (blue) nuclear counterstain. Scale bar 50μm. Images were obtained 72h following nanoneedles interfacing. (E) Immunofluorescence microscopy of Caspase 3/7 activation. Lack of nuclear staining with faint cytoplasmic staining 72h following nanoneedle interfacing (nN), comparable to untreated control (ctr) demonstrate lack of Caspase 3/7 activation, indicating absence of apoptotic events. Caspase 3/7 (green) staining with DAPI (blue) nuclear counterstain. Scale bar 50μm.

These data indicate that nanoneedles interfacing with primary HCEnCs *in vitro* by centrifugation retains cell morphology and ZO-1 expression, and does not induce apoptosis.

### Targeted silencing of p16 in HCEnCs in vitro

We then determined the efficiency of siRNA nanoinjection and its ability to induce targeted gene silencing *in vitro*. Microscopy analysis revealed fluorescently-labeled siGlo Red siRNA abundantly and uniformly loaded onto the nanoneedles prior to interfacing with the cells in culture (Figure 2A). Following nanoinjection, the siGlo was delivered to the cytoplasm of HCEnCs, in 27.6±8% of the treated cells, as quantified from immunofluorescence images (Figure 2B). Three strains of primary HCEnCs derived from different donors were used to assess p16 silencing upon nanoinjection. The delivery of p16-targeting siRNA to primary HCEnCs resulted in a significant (*p*=0.04) silencing of the target gene by 23±7% with respect to the non-specific control (NSC) (Figure 2C). When normalised for the 27.6% transfection efficiency, this approach effectively yielded a 72.4±3.5% silencing of p16 within transfected cells. These results demonstrate the transfection of primary HCEnCs *in vitro* by siRNA nanoinjection, resulting in significant silencing of the target p16 gene in transfected cells.

**Figure 2.**
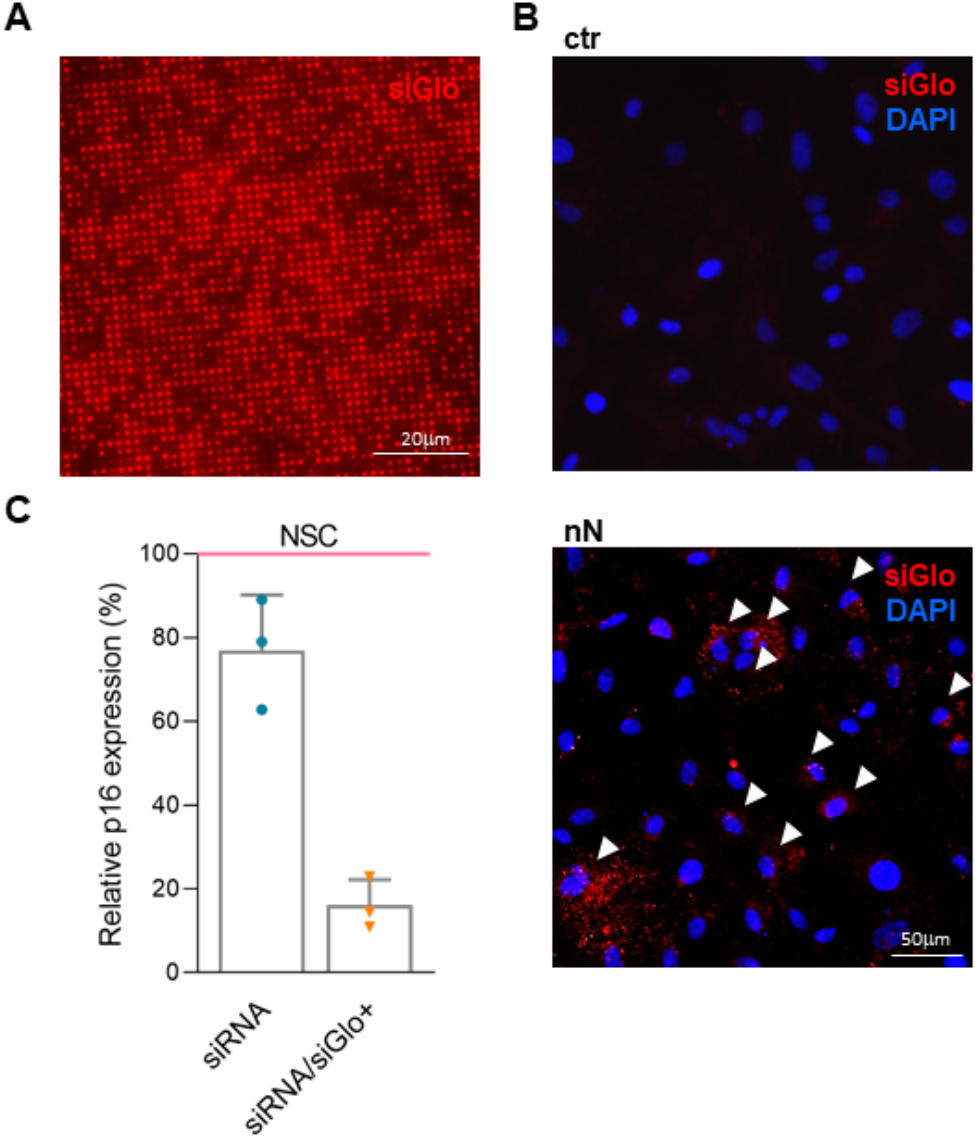
*In vitro* nanoinjection of p16 (CDKN2A) siRNA in HCEnCs. (A) Fluorescence microscopy of nanoneedles loaded with siGlo siRNA. siRNA is adsorbed uniformly across the nanoneedles. Scale bar 20μm. (B) Fluorescence microscopy of HCEnCs 48h following nanoinjection of siGlo. siRNA accumulates in the cytosol of the cells upon nanoinjection (nN), as compared with the untreated HCEnCs (ctr). White arrows indicate some of the highly transfected cells. siGlo signal (red) with DAPI (blue) nuclear counterstain. Scale bar 50μm. (C) RT-PCR of p16 expression showing silencing 48h following p16-siRNA nanoinjection, normalised and compared to NSC (non-specific control, pink line). Experiment performed on three primary HCEnCs strains derived from different donors at passage 1 in culture. The bar on the left (dark blue) indicates overall silencing level, the bar on the right (light blue) is normalised to the fraction of siGlo-transfected cells in culture.

### Effects of nanoinjection in vitro

We then evaluated the functional effects of p16 silencing in HCEnCs *in vitro*. Immunofluorescence quantification of p16-expressing cells revealed that p16 siRNA nanoinjection downregulated target protein expression when compared with NSC (Figure 3 A and B). The fraction of p16-expressing cells decreased from 57±13% for NSC nanoinjection to 35.2±3.8% for p16 siRNA-nanoinjection (Figure 3B). This difference is statistically significant (*p*=0.002) and represents a 21.8% reduction in p16 expressing cells, which aligns well with the observed 27% delivery efficiency and the 23% overall p16 silencing (Figure 2). When normalising the 21.8% reduction in p16 expressing cells for the 27.6% transfection efficiency, this approach effectively yielded a 79% knockdown. These data indicate that the primary human cells, which are transfected through nanoinjection, effectively silence target gene expression and knockdown the synthesis of the target protein.

**Figure 3.**
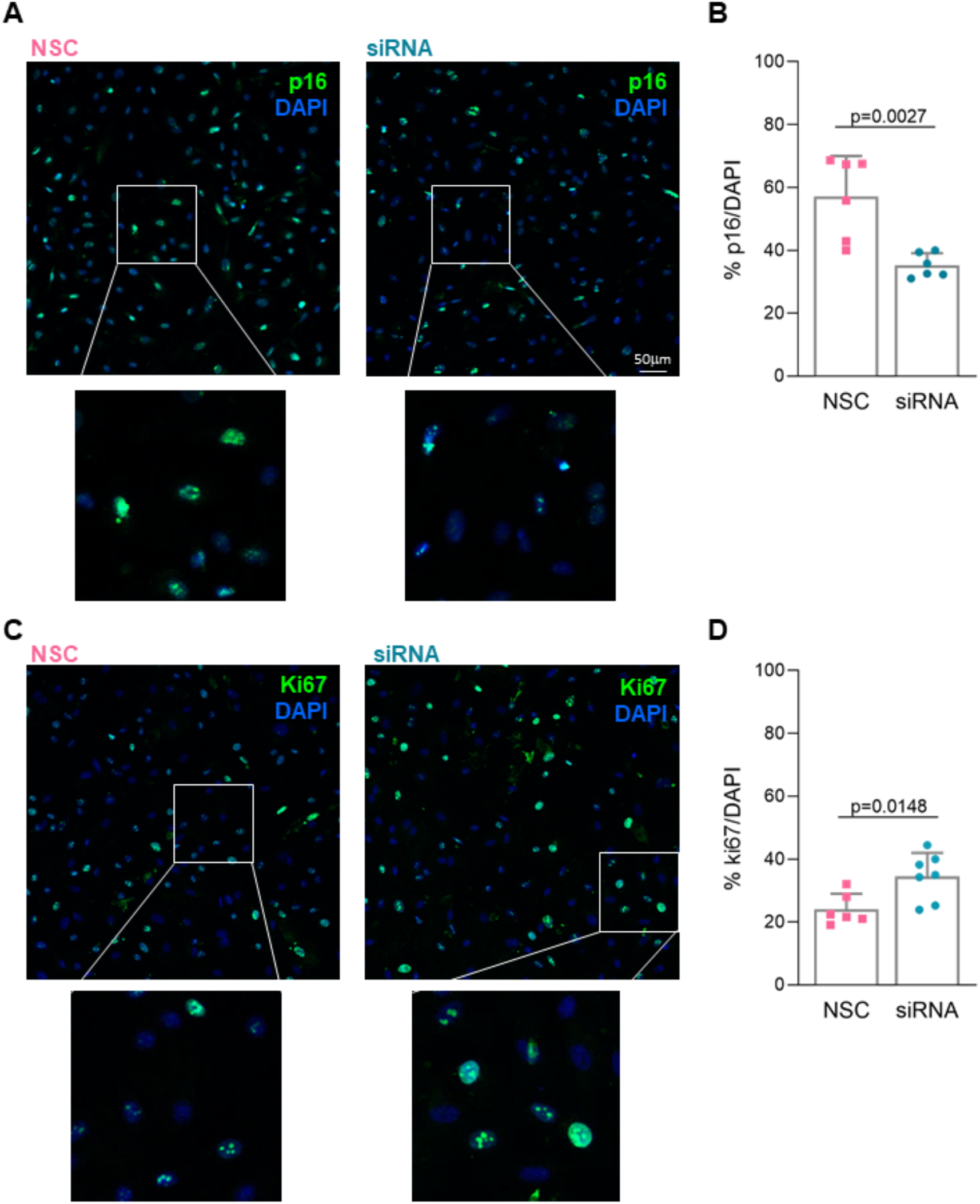
Effects of *in vitro* nanoinjection. (A,B) Nanoinjection with p16-siRNA (siRNA) induces p16 protein knockdown if compared to NSC nanoinjection (NSC) 72h following interfacing. (A) Immunofluorescence microscopy showing p16 protein expression. p16 staining (green) with DAPI (blue) nuclear counterstain. Scale bar 50μm. (B) Quantification of the fraction of cells expressing p16 for p16-siRNA (siRNA) and NSC nanoinjected HCEnCs. p16-siRNA nanoinjected samples have a statistically significant lower fraction of p16-positive cells. (C,D) RNAi nanoinjection to HCEnCs enhances their proliferative capacity 72h following interfacing. (C) Immunofluorescence microscopy showing ki67 protein expression in p16-siRNA (siRNA) and NSC treated HCEnCs *in vitro*. ki67 staining (green) with DAPI (blue) as nuclear counterstain. (D) Quantification of the fraction of cells expressing ki67 protein in p16-siRNA (siRNA) and NSC treated HCEnCs *in vitro*. p16-siRNA nanoinjected samples have a statistically significant higher fraction of ki67-positive cells.

The reduction of p16 protein expression was supported by a concomitant upregulation of ki67, indicative of an increased proliferation in p16 siRNA-nanoinjected cells (Figure 3C and D). The fraction of ki67 positive cells following siRNA nanoinjection was 34.4±7.5%, a significant (p=0.014) increase of 10.4% with respect to the 24.0±4.9% observed for NSC nanoinjection (Figure 3D). These data indicate that RNAi nanoinjection therapy has functional outcomes that impact the desired pathway regulation in primary somatic human cells *in vitro*.

### Nanoneedle interfacing with the human corneal endothelium

The successful nanoinjection of HCEnCs *in vitro* encouraged us to develop a nanoinjection approach for the endothelium in whole corneas (Figure 4). Several delivery studies have exploited *ex vivo* explanted human corneas with upper sided endothelial surface^27,28^: the availability of corneas deemed unsuitable for transplantation and discarded from eye banks^2^ make those tissues a precious resource for preliminary studies. Moreover, the 5μm thickness of the corneal endothelium monolayer well matches the length of the nanoneedles, insuring that all HCEnCs can be targeted simultaneously while preventing reaching other cells in the underlying tissues^25^. For nanoinjection in explanted corneas, nanoneedle chips loaded with siRNA were placed on the endothelial side of human cornea explants and interfaced with the assistance of centrifugation (Figure 4A). Confocal microscopy shows multiple nanoneedles co-localising with the cytosol and nucleus of cells throughout the endothelial layer, indicating successful interfacing (Figure 4B). A 3D reconstruction allowed visualizing the nanoneedle array through, but not beyond, the cells thickness (Figure 4C), which is confirmed by the orthogonal projections (Figure 4D). The cells retained the expected expression and localization of ZO-1 at 72h post-nanoinjection, indicating that HCEnCs maintained their morphological integrity and ZO-1 marker expression across the whole endothelial monolayer upon nanoneedle interfacing (Figure 4E). Similarly to what observed *in vitro*, nucleic acids nanoinjection also preserved cell viability, as shown by the absence of apoptosis events at 72h (Figure 4F). DNA loaded within the nanoneedles (Supp Figure 1A) can be further nanoinjected to the cytoplasm of HCEnCs *ex vivo* (Supp. Figure 1B) and a 3D reconstruction helps visualizing cytosolic distribution of the nucleic acids following nanoinjection (Supp. Figure 1C).

**Figure 4.**
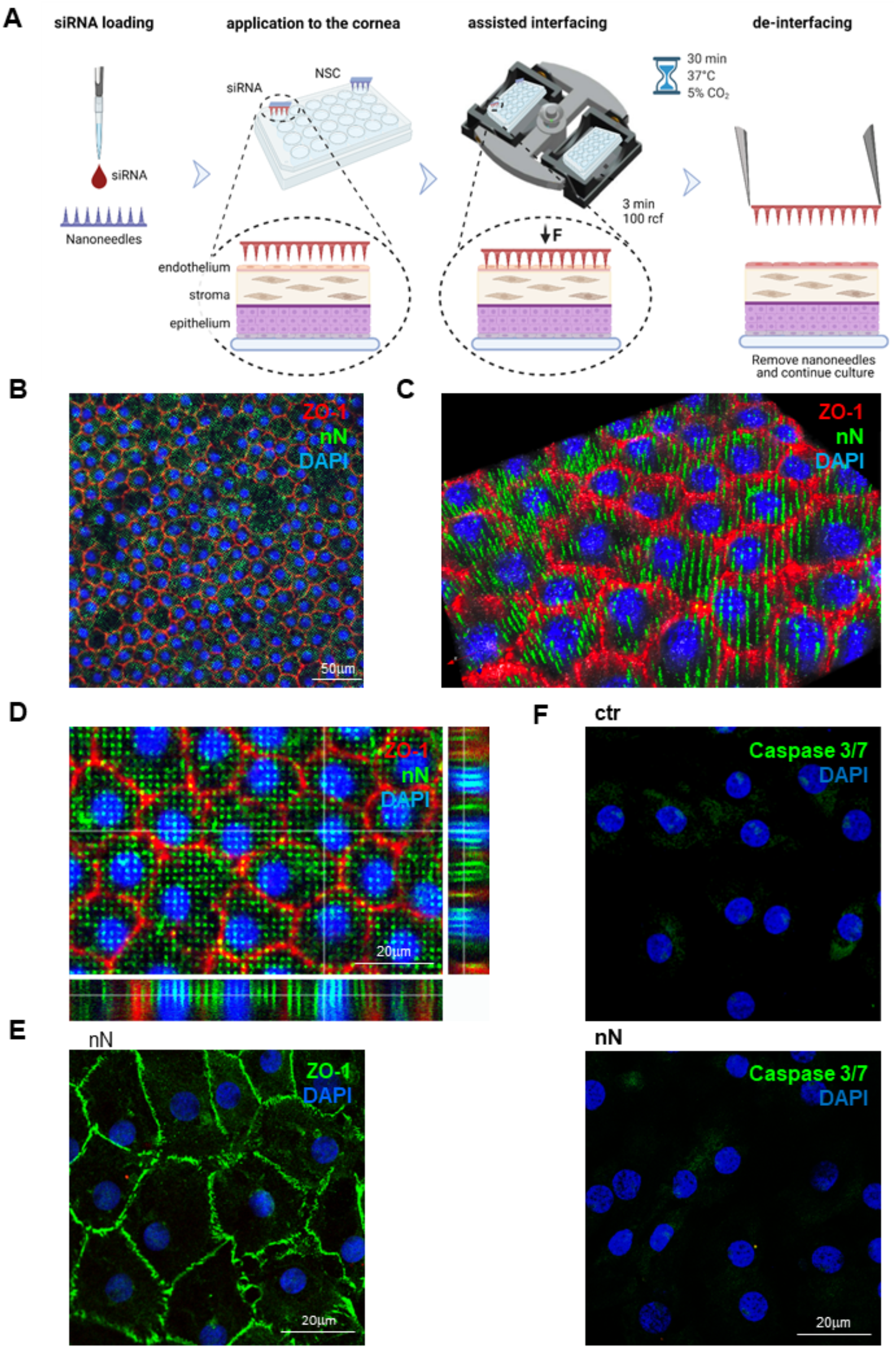
Nanoneedle interfacing with the explanted human corneal endothelium. (A) Schematic representation of the nanoinjection approach for explanted human corneas. Image created with Biorender.com (B-D) Immunofluorescence confocal microscopy of the interface between nanoneedles and the endothelium of human cornea explants. Images were obtained immediately after nanoneedle assisted interfacing by centrifugation. Nanoneedles co-localised with HCEnCs and did not protrude beyond them. ZO-1 (red) localizes in HCEnCs membrane, FITC (green) labels nN and DAPI (blue) nuclear counterstain. (B) Large-scale overview of a single z-plane across the endothelium. Scale bar 50μm. (C) 3D reconstruction from Z-stack. (D) Orthogonal projections. Scale bar 20μm. (E) Immunofluorescence confocal microscopy image of the explanted corneal endothelium obtained 72h after nanoneedles interfacing, showing a maintained native endothelial morphology. ZO-1 (green) staining with DAPI (blue) nuclear counterstain. Scale bar 20μm. (F) Immunofluorescence microscopy of Caspase 3/7 activation 72h after nanoneedles interfacing. Lack of nuclear staining with faint cytoplasmic staining, comparable to untreated control demonstrate lack of Caspase 3/7 activation, indicating absence of apoptotic events. Caspase 3/7 (green) staining with DAPI (blue) nuclear counterstain. Scale bar 20μm.

These data indicate that the nanoinjection of corneal endothelial cells in human explanted corneas is feasible and non-toxic.

### Effects of nanoinjection to the human corneal endothelium

We assessed the effects of nanoinjection for RNAi therapy in the endothelium of human corneal explants (Figure 5). A mix of siGlo and p16 siRNA were loaded on the nanoneedles and interfaced with the corneal endothelium. Nanoinjection delivered the siRNA into the cytoplasm of HCEnCs within the endothelial layer as visualised through the siGlo fluorescence (Figure 5A). Within untreated corneal endothelial layers, cells were uniformly p16 positive (Figure 5B), indicative of the corneal endothelium proliferative block. Upon nanoinjection, siRNA transfection induced p16 knockdown (Figure 5C-D). In the selected area of interfacing, 10±0.7% of HCEnCs were transfected with siGlo. Overall, 12.2±1.2% of cells showed a downregulation of p16 protein, including 80.4±2.9% of the siGlo-transfected cells and 5±1.1% of the non siGlo-transfected cells (Figure 5D). The high-magnification inserts further show the strong correlation between HCEnCs siGlo transfection (red signal) and p16 protein knockdown (green), as indicated by the asterisks. The p16 signal intensity significantly (p=1.5^-8^) decreased in siGlo transfected (siGlo+, 18±21.5%), as compared to untransfected cells (siGlo-, 100±30%), confirming that RNA therapy through nanoinjection is effective at silencing p16 (Figure 5E).

**Figure 5.**
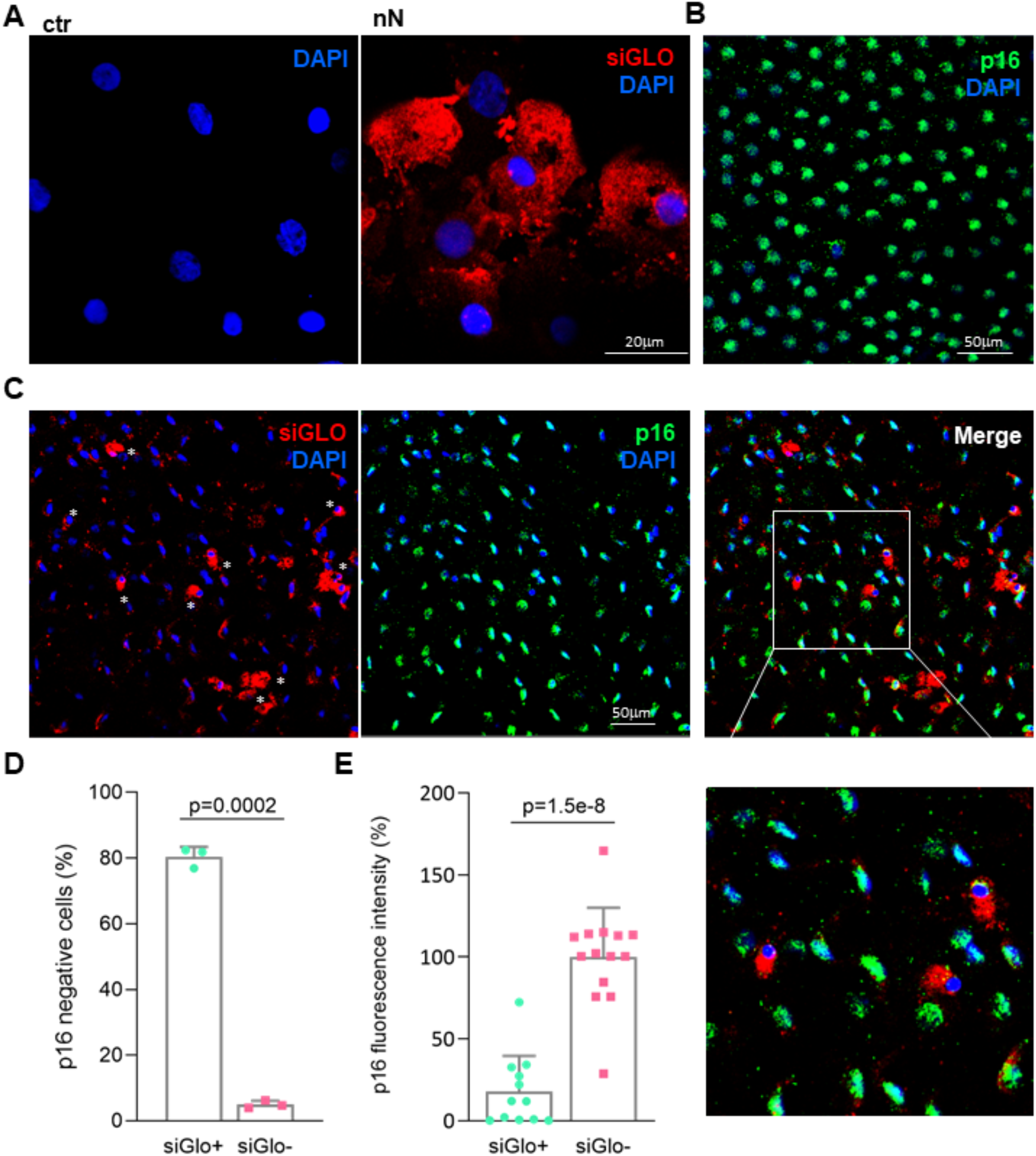
Effects of p16-siRNA nanoinjection to the explanted human corneal endothelium. (A) Immunofluorescence microscopy of siGlo nanoinjection to the endothelium of explanted human corneas. Images were obtained 48h after nanoinjection. HCEnCs cells display cytosolic siGLO in the area of nanoinjection (nN) as compared to untreated controls (ctr). siGlo signal (red) with DAPI (blue) nuclear counterstain. Scale bar 20μm. (B) Immunofluorescence microscopy of p16 protein expression in the untreated endothelium of explanted corneas. Nuclear expression of p16 can be detected in almost all HCEnC of the cornea after 72h of *ex vivo* culture. p16 (green) staining with DAPI (blue) nuclear counterstain. Scale bar 50μm. (C) Immunofluorescence microscopy of explanted human corneas 72h following nanoinjection of p16 siRNA. A significant correlation is visible between siGlo signal and loss of p16 signal, as highlighted by the white asterisks. p16 (green), siGlo (red) staining with DAPI (blue) nuclear counterstain. Scale bar 50μm. (D) Immunofluorescence quantification evaluating the fraction of p16 negative cells in siGlo+ transfected and siGlo-untransfected HCEnCs. (E) Immunofluorescence quantification of p16 expression levels in siGlo+ transfected and siGlo-untransfected cells.

## Discussion

Nanoneedles can efficiently deliver several types of nucleic acids to hard-to-transfect cells, among which primary neural, immune and stem cells without appreciably altering their phenotype^23^. This study shows that nanoinjection can mediate RNAi in human tissues and provide therapeutically-relevant outcomes including inducing desired changes in cell function.

Nanoinjection could transfect primary HCEnCs and the endothelium of a human cornea with minimal cellular invasiveness: *in vitro* or *ex vivo* nanoinjected HCEnCs maintained cell morphology and ZO-1 expression; cells appeared functional without signs of apoptosis (Figure 1 and 4), which is an important feature in corneal quality provided a 72% silencing of p16 mRNA in transfected cells (Figure 2) which knocked down p16 protein expression and induced the desired functional effect of increased HCEnCs proliferation (Figure 3). This enhanced proliferation can be leveraged to increase HCEnCs density, fundamental for improving donor corneas availability and for advanced therapies based on *in vivo* transient induction of HCEnCs proliferation or *in vitro* cell expansion.

Nanoinjection to explanted corneas showed interfacing throughout the limited thickness of the corneal endothelium (Figure 4A-C and Supp. Figure 1). The p16 knockdown was observed in more than 80% of the siGlo-transfected HCEnCs (Figure 5), highlighting the potential to further develop nanoinjection for *in vivo* corneal endothelium reprogramming with minimal mechanical invasiveness. The 80% knockdown in p16 protein expression for the siGlo-transfected cells matched the 79% p16 protein knockdown within transfected cells observed *in vitro* (Figure 3).

Nanoinjection provides efficient, non-immunogenic transfection for different tissues^23^, which was confirmed herein for the non-mitogenic corneal endothelium. Future integration of a flexible nanoneedle substrate to conform to the target tissue alongside optimization of nanoneedle topography and surface chemistry to match cell requirements^21,23^ should provide desirable enhancements of transfection efficiency. These optimization steps would also improve the interfacing with the inner surface of the intact cornea, enabling integration of this approach within current corneal endothelium surgery techniques with minor adaptations for *in vivo* corneal endothelial gene therapy.

In conclusion, this study assessed the feasibility of nanoinjection for RNAi therapy in human endothelial corneal cells in culture and in explanted human corneas, demonstrating targeted siRNA transfection, gene silencing, protein knockdown and functional outcomes.

## Materials and Methods

### Fabrication of the nanoneedles

Porous silicon nanoneedles are fabricated according to our established protocols^29,30^ over the entire surface of a 100mm, <100>, p-type silicon wafer. First a 160nm layer of silicon-rich silicon nitride is deposited by chemical vapour deposition. The substrate is then patterned by UV photolithography with a square array of 600-nm diameter dots with 2μm pitch. For the photolithography a 220nm layer of NR9-250P photoresist (Futurrex Inc, USA) is spin coated on the substrate with the following parameters 500RPM/1000RPMS/5s, 4000RPM/5000RPMS/40s. The substrate is pre-baked at 70C for 180s on a hotplate followed by hard vacuum contact exposure in an MA6 mask aligner (K. Suss GMBH, Germany). The exposed substrate is post-baked for 60s at 100C on a hotplate, developed in 3:1 RD6:H_2_O developer solution for 12s (Futurrex Inc, USA) rinsed with excess water and dried with N_2_. The photolithographic pattern is transferred into the silicon nitride layer by reactive ion etching (Oxford Instruments, NGP80) with the following parameters: 50 sccm CHF_3_, 5 sccm O_2_, 150 W forward power, 55 mTorr pressure, 150 s. The remaining photoresist was stripped with acetone and the substrated cleaned with isopropanol and dried under nitrogen stream. For metal assisted chemical etching (MACE) the native oxide layer was first removed by dipping the substrate in 10% hydrofluoric acid (HF, Honeywell, USA) for 2 minutes, immediately followed by electroless Ag deposition for 2 minutes in 100ml of 20 mM AgNO_3_ (Sigma Aldrich) in 10% HF. The substrate was rinsed in water and isopropanol and dried under nitrogen stream. The MACE process formed the porous silicon pillar structures by dipping the substrate in 400ml of a solution composed of 1 part 30vol H_2_O_2_ (Sigma Aldrich) and 99 parts 10% HF solution. To stop the etch, the wafer was dipped in DI water, then rinsed with excess water and isopropanol, and dried under nitrogen stream. The residual Ag was removed in gold etchant solution (Aldrich) for 10 minutes. The substrate was rinsed with excess water, isopropanol and dried under nitrogen stream. The final conical nanoneedle structure was obtained by reactive ion etching in an NGP80 (Oxford Instruments, UK) in the following conditions: 20 sccm SF_6_, 300W, 100 mTorr for 120s. The 100mm substrate was diced in 8×8mm chips (Disco, DAD3220) for use in 24-well plates. Individual chips were oxidised prior to use by oxygen plasma in a Femto plasma asher (Diener, Germany) at 10sccm O_2_, 0.2 mBar, 100W for 10 minutes.

Fluorescently-labelled nanoneedles were obtained by conjugating fluorescein isothiocyanate (FITC) to the oxidised silicon nanoneedles through a silane linker. Aminopropyltriethoxysilane (APTES) was grafted on the silicon surface in 2% APTES ethanoic solution for 2h. The substrate was then washed 3 times in ethanol and 1 time in DI water. The APTES-functionalised nanoneedles were reacted in a phosphate-buffered solution (PBS) of 0.01 mg/ml FITC for 1h. The substrate was washed 3 times in PBS and 1 time in DI water, and dried under nitrogen stream until further use.

### Ethical statement

Human donor corneas, unsuitable for transplantation, were procured by Italian Eye Banks after obtaining written consent from the donor’s next of kin for research use. The experimental protocol was approved by ISS-CNT (Italian National Transplant Centre) and by the local ethical committee (Comitato Etico dell’Area Vasta Emilia Nord, p. 0002956/20). The tissues were handled in accordance with the declaration of Helsinki.

### Nanoinjection of human corneal endothelial cells in vitro

Human corneas, preserved in Eusol at 4 °C, were selected for experiments with the following criteria: age ranging from 4 to 90 years old, no history of corneal diseases, HCEnCs density greater than 1,800 cells/mm^2^, death to preservation interval lower than 15 h and used for cultures within 15 days from death (Table 1). The peel and digest method was used to obtain primary culture of HCEnCs. Briefly, intact Descemet’s membrane was stripped off the corneas and HCEnCs isolated using 1.5mg/ml Collagenase A (Roche, USA) in DMEM (Thermo Fisher Scientific, USA) for 2h at 37 °C. Isolated HCEnCs were then pelleted at 1,200 rpm for 3 min. A further dissociation step with TrypLE (Thermo Fisher Scientific, USA) for 5 min at 37 °C helped cell-cell isolation. After that, the cells were pelleted at 1,200 rpm for 3 min and plated in 24 well plates coated with FNC Coating Mix (AthenaES, USA). Dual media method^31^ was used for expansion of HCEnCs, which were cultured at 37 °C in 5% CO_2_, and the medium was changed every 2 days. HCEnCs between the first and the third passages were employed for experiments: 2×10^4^ cells were plated in a chambered 8 well coverslips (IBIDI) 24h before being treated with nanoneedle chips.

**Table 1.** List of donor human corneas used for the experiments.

| Cornea n. | sex | age | D/P | Endo count | experiment |
| --- | --- | --- | --- | --- | --- |
| 1 | M | 80 | 21.4 | 2500 | Delivery method optimization |
| 2 | M | 51 | 2.1 | 2700 | Delivery method optimization |
| 3 | F | 81 | 4.2 | 2500 | Method optimization + Caspase |
| 4 | M | 84 | 4 | 2500 | Method optimization + Caspase |
| 5 | M | 63 | 12 | 2600 | Method optimization |
| 6 | M | 90 | 4.55 | 2600 | Method optimization |
| 7 | M | 79 | 13 | 2700 | Method optimization plasmid |
| 8 | F | 88 | 2.2 | 2800 | Method optimization plasmid |
| 9 | F | 60 | 20 | 3100 | Ex vivo plasmid |
| 10 | F | 77 | 7.2 | 2500 | Ex vivo siGlo |
| 11 | F | 82 | 17.5 | 2800 | Ex vivo siGlo |
| 12 | F | 76 | 8 | 2300 | Ex vivo plasmid |
| 13 | F | 85 |  |  | Ex vivo siGlo + plasmid |
| 14 | M | 79 |  |  | Ex vivo siGlo + plasmid |
| 15 | F | 80 | 10 | 2217 | Ex vivo p16 Ab optimization |
| 16 | M | 83 | 5.2 | 1501 | Ex vivo p16 Ab optimization |
| 17 | M | 78 | 10 | 2057 | Ex vivo p16 Ab optimization |
| 18 | F | 72 | 21 | 1639 | Ex vivo siRNA delivery |
| 19 | F | 79 | 23 | 2283 | Ex vivo siRNA delivery |
| 20 |  | 81 | 15.3 | 2057 | Ex vivo siGlo delivery + siRNA KD |
| 21 |  | 81 | 15.3 | 2222 | Ex vivo siGlo delivery + siRNA KD |
| 22 |  | 80 | 21.5 | 2192 | Ex vivo siGlo delivery + siRNA KD |
| 23 |  | 80 | 21.5 | 2120 | Ex vivo siGlo delivery + siRNA KD |
| 24 | M | 67 | 6 | 1900 | In vitro siGlo delivery + siRNA KD |
| 25 | F | 77 | 23 | 1700 | In vitro siGlo delivery + siRNA KD |
| 26 | M | 45 | 10 |  | In vitro siGlo delivery + siRNA KD |
| 27 | F | 78 | 20 |  | In vitro siGlo delivery + siRNA KD |
| 28 | M | 66 | 21 | 1800 | In vitro siGlo delivery + siRNA KD |
| 29 | F | 48 | 20 |  | Ex vivo interface ZO-1 |
| 30 |  | 26 | 21 | 3268 | In vitro Caspase and interface |

Nanoneedle treatments starts by placing the chips (8 × 8 mm) at the bottom of a 24 well plate, washing them with 2M HCl to remove impurities and then rinsing twice in distilled water. siRNAs (100nM) were then loaded onto the chip in a total volume of 10ul, dissolved in a buffer composed of 0.25M Glycine and 400mM KCl, pH 5, and incubated for 30min. The chip was then applied facing down over the cells monolayer where medium was removed and spun at 350 rcf for 3minutes in a swinging bucket centrifuge. Fresh medium was added to the well and the chip was removed after 30 minutes of incubation.

siRNAs loaded onto nanoneedle chip for the experiments were: siGloRed Transfection Indicator (Dharmacon), Silencer Select Validated siRNA CDKN2A (s218, Thermo Fisher) and Silencer™ Select Negative Control (4390843, Thermo Fisher), indicated as Non Specific Control (NSC). Plasmid DNA (pm-mCherry-N1, Addgene) was labelled with Label It DNA kit (Mirus), following the manufacturer’s instructions.

### Nanoinjection of explanted human corneas

Human corneas, preserved in Eusol at 4 °C, were used for experiments within 15 days from explant. Nanoneedle chips (8 × 8 mm) were loaded with siRNAs (100nM) as described above in the Nanoinjection *in vitro* section.

Corneal buttons were removed from Eusol, washed in DPBS, cut in quarters and each one was located into a well of a 24well plate with the corneal endothelium facing up. The chip was placed facing down onto the top of the cornea, in direct contact with corneal endothelium, and the plate was then spun at 100 rcf for 3 minutes in a swinging bucket centrifuge. The chip was left in contact with the cells for further 30 min of incubation with DMEM (Thermo Fisher Scientific, USA), 4% fetal bovine serum (FBS, Fisher Scientific, USA), 4% dextran (Sigma Aldrich, USA), penicillin/streptomycin (Euroclone, Italy) at 37°C and 5% CO_2_ and then placed in a new well, faced up, with fresh medium.

### RT-PCR

RNeasy plus Micro Kit (Qiagen) was used to extract RNA from HCEnCs, which was then quantified through the Nanodrop 100 (Thermo Fisher Scientific) and reverse transcribed into cDNA with the High Capacity cDNA Reverse Transcription Kit (Thermo Fisher Scientific). RT-PCR assays were performed using 7900HT Fast Real-Time PCR System (Thermo Fisher Scientific), choosing the following TaqMan Real Time PCR Assays probes: Human CDKN2A (Hs00923894_m1) and Human GAPDH (Hs02786624_g1). ΔCt and ΔΔCt calculations using GAPDH as housekeeping control were performed to evaluate effective RNA expression. For each condition, all complementary cDNA samples were run in triplicate. Human primary corneal endothelial cultures isolated from three different subjects were used at passage 1 for RT-PCR analysis.

### Immunofluorescence and whole mount imaging

Samples were washed in PBS and fixed in 3% paraformaldehyde (PFA) for 15 minutes at room temperature (RT) and permeabilized by 0.5% Triton x-100 (Bio-Rad, USA) for 10 minutes. A blocking solution composed of 2% bovine serum albumin (BSA; Sigma-Aldrich, USA), 2% FBS and 0.01% Triton in PBS was used to saturate the non-specific binding sites for 30 min at 37 °C. Primary and secondary antibodies were diluted in blocking solution and incubated for 1 h at 37 °C. Nuclei were counterstained with DAPI (1:40.000 dilution, Roche, USA) for 5 min at RT. Three rinses in BSA 0.2% were performed between all steps, except before incubation with primary antibody. The corneal slice was finally placed on a glass slide with DAKO mounting medium (Agilent, USA), flattened using a glass coverslip and retained by adhesive tape.

The primary antibodies used were ZO-1 (1:100, 40-2200, Thermo Fisher USA), p16 (1:50, ab108349, abcam, USA), ki67 (1:100, ab15580, abcam, USA), while the secondary antibodies were Alexa Fluor 488 anti-rabbit, 1:2000, and Alexa Fluor 568 anti-mouse, 1:1000 (Thermo Fisher, USA). Quantification of p16 and ki67 staining was obtained counting the number of positive cells (primary antibody signal), relative to the total number of cells in that field (DAPI staining), expressed in percentage with standard deviation (3 fields for each replicate were collected). p16 fluorescence intensity (Figure 5) was evaluated using ImageJ software.

Cell apoptosis was evaluated with CellEvent® Caspase 3/7 Green (Thermo Fisher, UK), following the manufacturer’s instructions. HCEnCs treated with 10mM H_2_O_2_ for 2h were used as a positive control for the assay (Supp. Figure 2A and B)^32^. DNA plasmid was labelled with Label IT® (Mirus, USA). Reagents used for immunofluorescence are listed in Supp. Table1. A confocal microscope (LSM900 Airyscan—Carl Zeiss) was used to obtain the images.

## Acknowledgements

CC would like to thank the European Research Council (ENBION Grant 759577) for providing financial support to this project. EM would like to acknowledge Prof. Guido Maria Macaluso for making his laboratory and equipment available for the experiments.

Thanks must go in particular to the patients who donated their organs for medical purposes or research.

