## Supplementary material for "Nanoneedles for targeted siRNA silencing of p16 in the Human Corneal Endothelium"

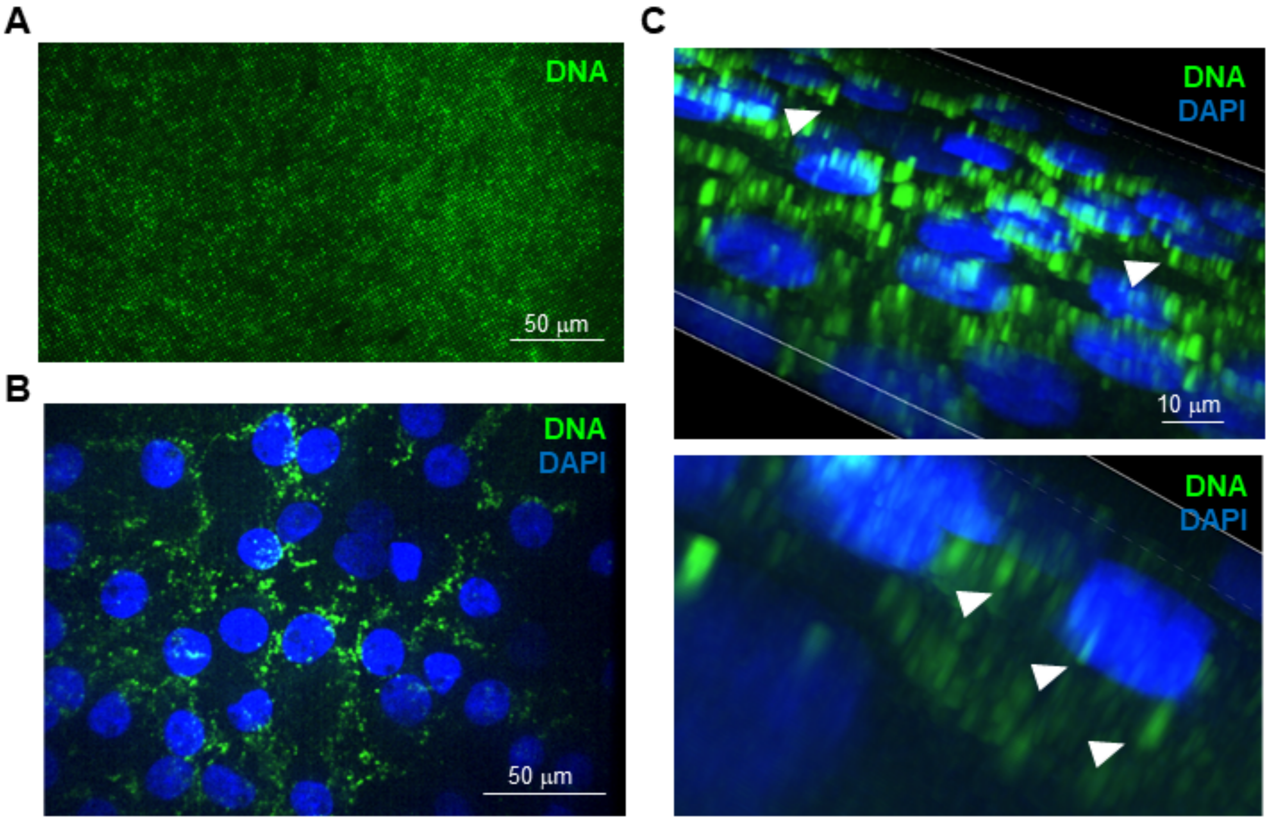


**Supplementary Fig1. Nucleic acid delivery to the endothelium of explanted human corneas.** (A) Fluorescence microscopy of nanoneedles loaded with an alexafluor-488-labeled DNA plasmid (green) before being applied to the human corneas. Scale bar 50μm. (B-C) Confocal Fluorescence microscopy of and explanted human cornea 24h following DNA plasmid (green) nanoinjection, showing cytosolic DNA signal within HCEnCs. DAPI (Blue) nuclear counterstain. (B) z-slice image. Scale bar 50μm. (C) 3D reconstruction. White arrows indicate cytoplasmic punctate signal within the HCEnCs. Scale bar 10μm.

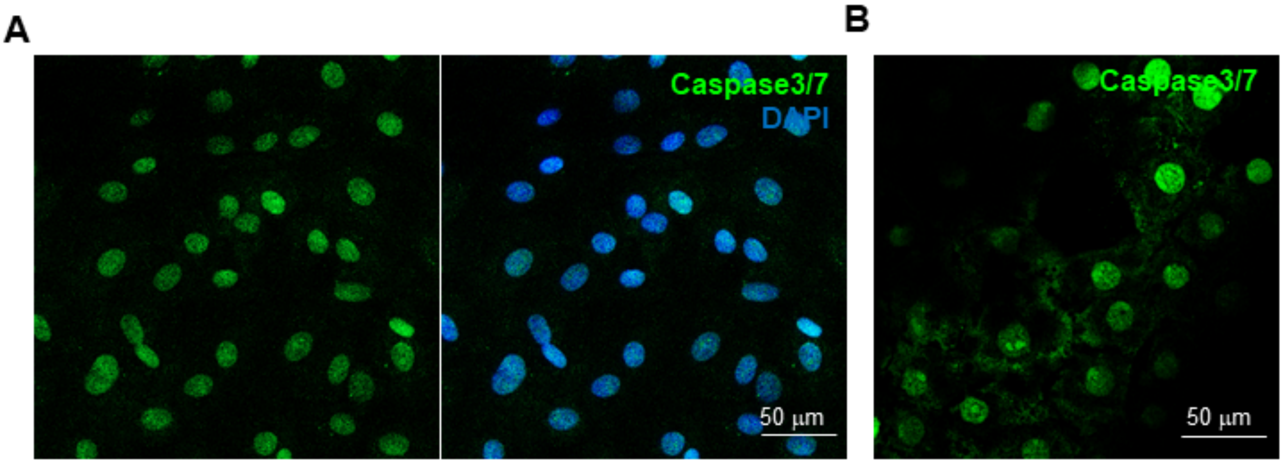


**Supplementary Fig.2 Caspase positive control in HCEnCs *in vitro* and *ex vivo*, p16 fluorescence intensity *ex vivo*** (A) Positive control for caspase activation assay in HCEnCs *in vitro.* Strong nuclear green signal indicates Caspase 3/7 activation and induction of apoptosis 2h following treatment with $H_2O_2$. DAPI (Blue) nuclear counterstain. Scale bar 50μm. (B) Positive control for caspase activation assay in explanted human cornea. Strong nuclear green signal indicates Caspase 3/7 activation and induction of apoptosis 2h following treatment with $H_2O_2$. Scale bar 50μm.

| Assay/protein | reference | dilution |
| --- | --- | --- |
| ZO-1 | 40-2200 (Thermo Fisher) | 1:100 |
| p16 | ab108349 (Abcam) | 1:50 |
| ki67 | ab15580 (Abcam) | 1:100 |
| CellEvent®Caspase 3/7 | C10723 (Thermo Fisher) | 1:200 |
| Lable It DNA kit | 3225 (Mirus) | |

**Supplementary Table1.** Antibodies and immunofluorescence reagents used for the experiments.
